# Behavioral Syndromes in Paper Wasps: links between social and non-social personality in *Polistes fuscatus*

**DOI:** 10.1101/2023.10.30.564788

**Authors:** Fatima W. Jomaa, Emily C. Laub, Elizabeth A. Tibbetts

## Abstract

Although much work has focused on non-social personality traits such as activity, exploration, and neophobia, there is a growing appreciation that social personality traits play an important role in group dynamics, disease transmission, and fitness, and that social personality traits may be linked to non-social personality traits. These relationships are important because behavioral syndromes, defined here as correlated behavioral phenotypes, can constrain evolutionary responses. However, the strength and direction of relationships between social and non-social personality traits remain unclear. In this project, we examine social and non-social personality traits, and the relationships between them, in the paper wasp *Polistes fuscatus*. With a novel assay we identify five personality traits, two non-social (exploration and activity) and three social (aggression, affiliation, and antennation) personality traits. We also find that social and non-social personality traits are phenotypically linked. We find a positive correlation between aggression and activity (r_s =_ 0.33) and a negative correlation between affiliation and activity (r_s =_ - 0.35). We also find a positive correlation between exploration and activity. Our work is an important step in understanding how phenotypic linkage between social and non-social behaviors may influence behavioral evolution. As a burgeoning model system for the study of genetic and neurobiological mechanisms of social behavior, *Polistes fuscatus* has potential to add to this work through exploring the causes and consequences of individual behavioral variation.

## Introduction

Animals exhibit consistent individual variation in a range of behaviors. Behavioral differences within a species that persist across different contexts are known as animal personalities or temperaments (Gosling 2008; Réale et al. 2007). Thus far, most research on animal personality has focused on non-social traits such as exploration, activity, and boldness (Bell, Hankison, and Laskowski 2009). Consistent non-social personalities have been identified in a wide range of vertebrates and invertebrates (Cote et al. 2011; Pinter-Wollman 2012). Less work has explored individual variation in social aspects of personality such as aggressive and affiliative behavior (Réale et al. 2007). However, there is growing evidence that many taxa have consistent individual variation in social personality, including unicellular organisms (Vogel et al. 2015), fish (Laskowski and Bell 2014), birds (Aplin et al. 2015), and mammals (Blumstein, Petelle, and Wey 2013).

Individual variation in social behaviors, forming social personalities or temperaments, is attracting increased interest due to wide ranging implications in ecological and evolutionary dynamics (Gartland et al. 2022). Social personality traits include sociability, aggression (Briffa, Sneddon, and Wilson 2015), cooperation (Sanderson et al. 2015), and have been observed in multiple taxa including birds (Aplin et al. 2013), mammals (Blaszczyk 2018; Taylor et al. 2012), and fish (Brask et al. 2019; Jacoby et al. 2014). Theory suggests social personality may have important effects on diverse behaviors including dispersal (Cote et al. 2010), foraging (Aplin et al. 2014), association formation (Cote, Fogarty, and Sih 2012), disease transmission (Drewe 2010), and collective behavior (Jandt et al. 2014; Jolles et al. 2015).

Social and non-social personality traits may form behavioral syndromes, where suites of personality traits are phenotypically correlated with each other (Sih, Bell, and Johnson 2004). Behavioral syndromes are hypothesized to arise when there is a common mechanism controlling multiple behaviors (e.g. hormonal pleiotropy, genetic linkage), correlational selection (Van Oers et al. 2004), or due to physiological allocation trade-offs (Veenema et al. 2003). For example, zebra fish that are more active also approach a predator dummy more often, consistent with an activity syndrome where underlying metabolic costs influence multiple types of active behavior (Moretz, Martins, and Robison 2007). Understanding whether there are consistent links between social and non-social behaviors is important because such links influence how traits respond to selection. For example, meta-analysis of additive genetic variance-covariance matrices suggest that behavioral syndromes may constrain potential evolutionary responses by an average of 33%, which is a larger constraint on selection than observed with life-history tradeoffs (Dochtermann and Dingemanse 2013). Therefore, assessing whether there are links between personality traits provides insight into both the mechanisms that produce animal personalities and how personalities respond to selection.

As interest in animal personality and behavioral syndromes has grown, so too has controversy over the best methodology to measure repeatability of behaviors and the strengths of relationships between them (Houslay and Wilson 2017; Wolak, Fairbairn, and Paulsen 2012; de Villemereuil et al. 2018). Identification of animal personality traits relies on determination that behaviors are ‘repeatable’, that individuals demonstrate more behavioral variation between individuals in population than within an individual (Bell, Hankison, and Laskowski 2009). However, how repeatability is calculated is highly variable, with many different statistical methods used, and disagreement about what experimental and individual variables should be included in the calculation of repeatability (Cauchoix et al. 2018; Evans et al. 2021; Uher, Asendorpf, and Call 2008). Including experimental variables may help control variation due to testing conditions, however including inappropriate variables may inflate repeatability by minimizing intraindividual variation due to other individual attributes. In addition, methods of determining behavioral syndromes are also controversial, with different approaches to handling within individual variation in multiple traits (Dingemanse, Dochtermann, and Wright 2010; Houslay et al. 2018; Mitchell and Houslay 2021). Further work is needed to detangle how differences in statistical analysis may impact detection of both personalities and behavioral syndromes.

*Polistes fuscatus* (*P. fuscatus*) paper wasps provide an interesting model system to explore behavioral syndromes because they exhibit significant variation in facultative cooperation. Nest-founding *P. fuscatus* queens can either start a nest alone or with a group of other cooperating queens (Roesler, 1991). *P. fuscatus* are also highly variable in the roles they perform on nest including foraging and defense – behaviors that have been linked to personality traits in other taxa (Walton and Toth 2016). Additionally, *Polistes* wasps are model organisms for studying facial recognition (Tibbetts 2004), dominance and reproductive skew in cooperative breeders (Jandt and Toth 2015; Reeve et al. 2000), and genomic underpinnings of cognition (Berens, Tibbetts, and Toth 2017), and chemical recognition (Cappa et al. 2020; Cini et al. 2019), all research areas that would benefit from incorporating personality data.

Here, we test *P. fuscatus* paper wasp nest-founding queens for the presence of repeatable variation in several commonly used social and non-social personality metrics. We also examine correlations between social and non-social personality traits to understand how social and non-social personality traits form behavioral syndromes. We assess two non-social behaviors (activity and exploration) and three social behaviors (aggression, affiliation, and anntenation behavior). Previous work has demonstrated that a close relative of *P. fuscatus, Polistes metricus* (*P. metricus*), demonstrates non-social personality traits (exploration and activity), and possibly social personality traits (aggression and boldness) (Wright et al. 2018). However, previous work measured aggression and boldness in response to predator intrusion, rather during conspecific interactions (Wright et al. 2017). This study uses a novel personality assay to evaluate social personality without interference from conspecific response, allowing us to examine the relationship between social and non-social personality traits. We also compare methods for assessing both personality repeatability and behavioral syndromes.

## Methods

### Wasp collection and care

Between May 6 -11 2021, we collected *P. fuscatus* foundresses (*n* = 74) from nine parks within a 30-mile radius of Minneapolis, Minnesota. Foundresses were collected after emerging from diapause and were all the approximately same age, having eclosed the previous August. Foundresses were collected before founding nests, or in the earliest stages of nest founding (fewer than 10 nest cells). During the early spring when they were collected, associations between foundresses are often ephemeral as they sample different nests and associations before forming stable social groups. Only two pairs of wasps were collected on a nest with another foundress. Wasps often disperse miles from where they eclosed from pupation, so wasps from a site are not highly related (Bluher, Miller, and Sheehan 2020). Foundresses were returned to the University of Michigan laboratory and stored individually in 4.5 x 3 in containers. Each wasp was fed sugar and water *ad libium*. None of the wasps died during testing.

### Behavioral assays

Behavioral assays took place between 9am and 3pm, which is the period when wasps are most active. Behavioral assays took place over a minimum 4-day period, although not all trials took place on a consecutive day-to-day basis due to constraints imposed by equipment availability, personnel, and to limit differences the in the time of day being tested. The maximum length of each assay testing period did not exceed 7 days. Wasps were given a minimum 24-hour respite period between within-assay trials. If an individual performed both affiliation and exploration trials on the same day (described below), the wasp was given a minimum hour respite period between trials. During recovery periods, wasps were returned to the environmental chamber in their original container to avoid further stimulation. Each foundress participated in 4 trials for each behavioral assay – trials lasted ten minutes and all trials were video recorded and later scored by three observers who were blind to experimental predictions.

### Dummy conspecific assay

To measure levels of affiliation, individual foundresses were placed inside a 10 x 10 x 2.5 cm lidded plexiglass compartment with an upright cardboard mounted dead dummy wasp for 10 minutes (Figure 1). Seven dummies were collected from the same populations as the focal wasps and were within the size range of typical wasps collected. Dummy wasps were freeze killed prior to trials and mounted in neutral body positions to standardize body posture. *Polistes* engage in typical aggressive and affiliative behaviors with freshly-killed conspecifics (Tibbetts, Vernier, and Jinn 2013). A separate dummy was used for each individual trial for each wasp to minimize dummy-specific effects on behavior. Between every trial, compartments were cleaned with 70% ethanol. Containers were left unlidded to dry for a minimum of 1 minute between trials to dissipate fumes.

**Figure 1:**
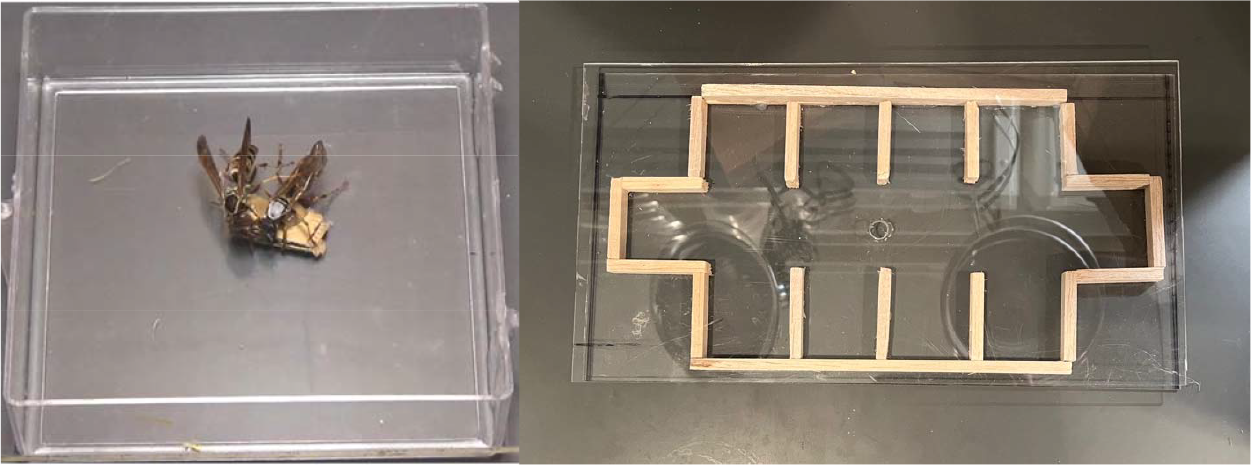
Arena setup for the dummy conspecific assay (left) and maze assay (right).

After videorecording trials for all individuals, we replayed each trial recording and scored behaviors. The following behaviors were observed: 1) bodily contact (wasp remains in stationary, non-aggressive contact with dummy), 2) antennation (wasp taps dummy with extended antennae, used in chemical assessment of other wasps, an “exploratory” social behavior), 3) dart (wasp lunges towards dummy), 4) dart with mandibles (wasp lunges towards dummy with mandibles open), 5) antenna drumming (wasp rapidly beats bent antennae on dummy wasp), 6) bite (wasp opens and closes mandibles on dummy), and 7) mount (wasp dominates dummy by climbing atop dummy and drumming antenna on the dummy’s head). Behavioral counts were then categorized into three categories: 1) affiliative (bodily contact), 2) antennation, and 3) aggressive (dart, dart with mandibles, antenna drumming, bite, mount). Aggressive and antennation behavior was totaled while affiliative behavior was recorded by time. Aggressive behaviors were log-transformed to normalize the data.

### Maze assay

The exploration assay was modeled off the assay used by Wright et al. (2018). To measure exploration and activity, wasps were placed into an ethanol cleaned 29 x 16 x 4 cm plexiglass arena with 10 built-in wooden chambers (Figure 1). A small hole was drilled into the plexiglass lid to allow entry. To conduct exploration trials, we allowed wasps to enter through the hole through placing their head and front legs into the hole and letting them crawl inside, then blockaded the hole with an additional square of plexiglass to prevent escape. Wasps were free to roam the arena for ten minutes. The number of individual chambers entered was counted and defined as exploration. Activity was defined as the duration of time the wasp spent actively moving inside the arena. Arenas were sprayed with 70% ethanol and allowed to dry for a minimum of 1 minute between trials.

### Analysis of behavioral repeatability

We calculated repeatability estimates for all behaviors using the RptR package in R, a package designed for analyzing repeatability by running a series of Linear mixed models (LMMs) for guassian distributed data and generalized linear mixed effects models (GLMMs) for poisson distributed data (R-core Team, 2023; Stoffel, Nakagawa, and Schielzeth 2017). Model fit was examined with residual plots generated with the performance package (Lüdecke et al., 2021). Aggression data was log-transformed before analysis to improve model fit. Aggression, activity, affiliation, and antennation were analyzed with guassian distributions and exploration was analyzed with poisson distribution. To compare differences in repeatability calculation based on variables included in estimates, we used two sets of models. To generate the most conservative estimates of repeatability, no fixed effects were included in the first set of “simple” models and individual ID was the only random effect included in the models. The second set of “adjusted” models included body mass and trial number as fixed effects. Activity and exploration “adjusted” models included individual id as a random effect and trial number and body weight as fixed effects. Affiliation, antennation, and aggression “adjusted” models included individual ID as a random effect and Dummy ID, focal wasp body weight, and trial number as fixed effects. The proportion of variation attributed to individual ID serve as our repeatability estimate, with a p-value cut-off of 0.05 for significance of the relationship between individual ID and personality measures along with confidence intervals excluding 0 to determine significant repeatability.

### Analysis of behavioral correlations

We used two different methods to examine correlations between behaviors. In the first analysis, data were averaged across trials for each individual before calculating correlations, then we assessed whether personality traits were linked by correlating personality traits using non-parametric Spearman rank correlations.

In the second analysis, we used bivariate mixed models to examine variation between behaviors while also retaining intraindividual variation in both compared behaviors. We fit generalized linear models (GLMMs) in a Baysian framework using Markov chain Monte Carlo techniques as outlined in (Houslay and Wilson 2017) with the package MCMCglmm (Hadfield 2010). We ran separate models to analyze covariance between each of our identified personality traits and included body mass and trail number as fixed effects for all personality traits and Dummy ID as a fixed effect for Affiliation, Aggression, and Antennation. Affiliation, Aggression, and Activity were analyzed with guassian distributions, while Antennation and Exploration were analyzed with poisson distributions. Gaussian distributed data were scaled to aid model fit. We estimated the mean covariance of each set of personality traits and the upper and lower bounds of the 95% credible interval of the covariance through creating posterior distributions of the among-individual correlation by dividing the corresponding individual covariance between each behavior by the product of the square root of their variances. Statistical significance of the correlation between two behaviors was determined by a 95% credible interval that did not include 0.

### Data Availability Statement

All data and code for analysis are available on the corresponding author’s GitHub profile (https://github.com/EmilyLaub/Wasp-behavioral-syndromes).

## Results

### Repeatability of Behavior

All 5 behaviors were significantly repeatable (exploration, activity, affiliation (non-aggressive body contact), aggression, and antennation (Table 1) when analyzed with both simple and adjusted models. Repeatability was calculated with only individual ID included in models (top line, straight text, Table 1) and with models that include additional individual and trial variables (lower line, italic, Table 1). We found that for Activity, Exploration, Activity, and Antennation, trial conditions and body mass (fixed effects) account for a relatively small proportion of variation in personality traits (less than 5%) and account for a smaller percentage of variation than Individual ID (Table 1, Table 2). However, trial conditions and body mass account for a greater proportion of variation than Individual ID for Aggression (Table 1, Table 2).

**Table 1:** Repeatability of measured personality traits. All values of R are statistically significant (P < 0.05). Values only individual ID included in models (top line, simple, straight text) and with models that include additional individual and trial variables (lower line, adjusted, italic).

| Behavioral Test | Personality Trait | R<br>(simple,<br>*adjusted*) | 95% CI<br>(simple, *adjusted*) | P-value<br>(simple, *adjusted*) |
| --- | --- | --- | --- | --- |
| **Maze** | Exploration | 0.497,<br>*0.513* | [0.343, 0.596],<br>*[0.357, 0.613]* | <0.001,<br>*<0.001* |
|  | Activity | 0.378,<br>*0.398* | [0.252, 0.505],<br>*[0.26, 0.504]* | <0.001,<br>*<0.001* |
| **Dummy**<br>**consppecific** | Affiliation | 0.344,<br>*0.377* | [0.22, 0.465],<br>*[0.253, 0.493]* | <0.001,<br>*<0.001* |
|  | Aggression | 0.141,<br>*0.159* | [0.031, 0.249],<br>*[0.037, 0.277]* | 0.003,<br>*0.001* |
|  | Antennation | 0.425,<br>*0.464* | [0.277, 0.54],<br>*[0.336, 0.592]* | <0.001<br>*<0.001* |

**Table 2:**
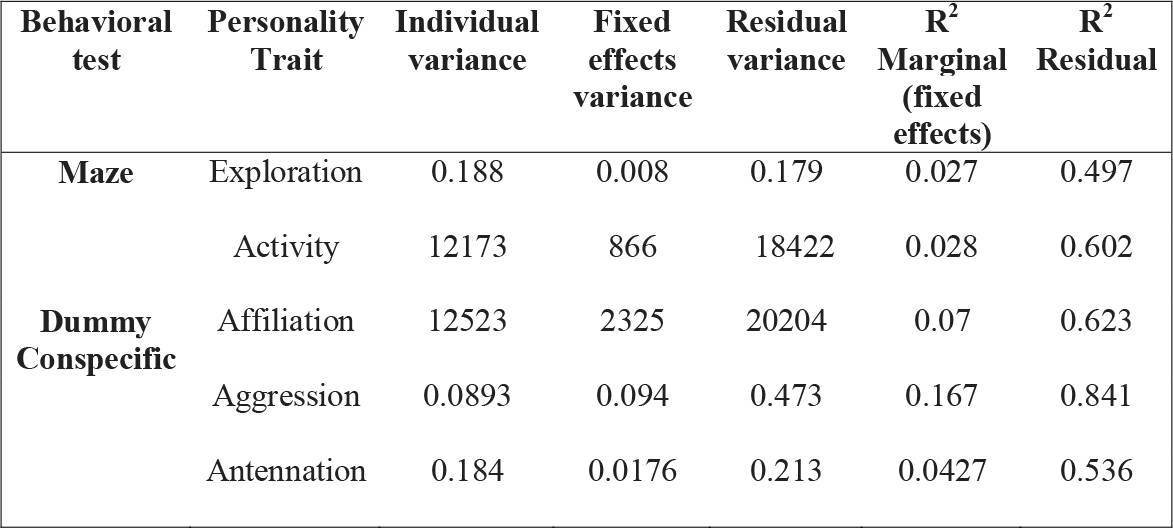
Variance and proportion of variation attributed to fixed effects and residuals for adjusted repeatability of each personality trait.

| Behavioral test | Personality Trait | Individual variance | Fixed effects variance | Residual variance | R<sup>2</sup> Marginal (fixed effects) | R<sup>2</sup> Residual |
| --- | --- | --- | --- | --- | --- | --- |
| **Maze** | Exploration | 0.188 | 0.008 | 0.179 | 0.027 | 0.497 |
|  | Activity | 12173 | 866 | 18422 | 0.028 | 0.602 |
| **Dummy**<br>**Consppecific** | Affiliation | 12523 | 2325 | 20204 | 0.07 | 0.623 |
|  | Aggression | 0.0893 | 0.094 | 0.473 | 0.167 | 0.841 |
|  | Antennation | 0.184 | 0.0176 | 0.213 | 0.0427 | 0.536 |

### Behavioral Correlations

In our first analysis, we assessed the relationship between personality traits using Spearman rank correlation after averaging behavioral scores across trials. We found that the two non-social personality traits, Exploration and Activity were linked. Wasps that exhibited greater explorative tendencies also displayed higher amounts of activity (*r*_S_ = 0.8883, *P* < 0.0001; Table 3)(Fig. 2a). However, there were no links between the three social personality traits (aggression, affiliation, antennation, (Table 2). Notably, there were some correlations between social and non-social personality. More active individuals were less affiliative than less active individuals (*r*_S_ = -0.3255, *P* = 0.0024; Table 3) (Fig. 2b). Activity was also significantly positively correlated with aggression (*r*_S_ = 0.331, *P* = 0.0036; Table 2) (Fig. 2c). There were no other significant correlations between social and non-social personality traits (Table 2).

**Table 3:** Spearman’s rank correlation coefficients (r_s_) and P-value (P) for correlations between wasp weight, exploration, activity, affiliation, and antennation. Bolded numbers indicate statistically significant correlations (p < 0.05)

|  | Exploration |  | Activity |  | Affiliation |  | Aggression |  | Antennation |  |
| --- | --- | --- | --- | --- | --- | --- | --- | --- | --- | --- |
| | $r_s$ | P | $r_s$ | P | $r_s$ | P | $r_s$ | P | $r_s$ | P |
| **Exploration** | - | - | **0.89** | **0.00** | -0.22 | 0.056 | 0.23 | 0.053 | -0.05 | 0.671 |
| **Activity** |  |  | - | - | **-0.35** | **0.002** | **0.33** | **0.003** | -0.02 | 0.849 |
| **Affiliation** |  |  |  |  | - | - | -0.1 | 0.415 | -0.09 | 0.455 |
| **Aggression** |  |  |  |  |  |  | - | - | 0.14 | 0.22 |
| **Antennation** |  |  |  |  |  |  |  |  | - | - |

**Figure 2:**
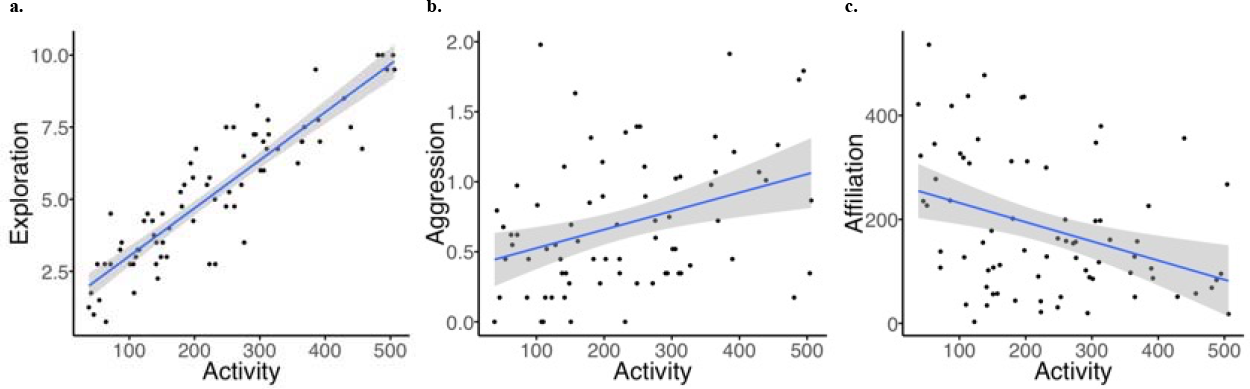
Significant phenotypic correlations from spearman’s rank correlations: a) Activity and Exploration (r_s_ = 0.89, p = 0.00), b) Activity and Aggression (r_s_ = 0.33, p = 0.003), c) Activity and Affiliation (r_s_ = -0.35, p = 0.002). Blue shading indicates standard error.

In our bivariate analysis, we assessed covariance between behaviors using GLMMs and Baysian framework using Markov chain Monte Carlo techniques. We found significant correlations between non-social personality traits with more active wasps also exploring more chambers (mean correlation: 0.943, 95% CI: [0.872, 1.00], Table 4, Fig. 2). We did not find any significant correlations between social personality traits (Table 4, Fig. 3). Excitingly, we found significant relationships between two non-social personality traits (Activity and Exploration) and two social personality traits (Aggression and Affiliation). Wasps that were more active were more aggressive (mean correlation: 0.623, 95% CI: [0.292, 0.956], Table 4, Fig. 3), but less affiliative (mean correlation: -0.481, 95% CI: [-0.753, -0.219], Table 4, Fig. 3). Wasps that were more exploratory were more aggressive (mean correlation: 0.462, 95% CI: [0.108, 0.865], Table 4, Fig. 3) but less affiliative (mean correlation: -0.352, 95% CI: [-0.599, -0.0413], Table 4, Fig. 3).

**Table 4:** Estimation of behavioral correlations from bivariate analysis. Five combinations of behaviors demonstrated significant correlations (bold): Activity and Exploration (mean correlation: 0.943, 95% CI: [0.872, 1.00]), Activity and Aggression (mean correlation: mean correlation: 0.623, 95% CI: [0.292, 0.956]), Activity and Affiliation (mean correlation: -0.481, 95% CI: [-0.753, -0.219]), Exploration and Affiliation (mean correlation: -0.352, 95% CI: [-0.599, -0.0413]), and Exploration and Aggression (mean correlation: 0.462, 95% CI: [0.108, 0.865]).

| Behavior combinations | Correlation | 95% Credible Interval |
| --- | --- | --- |
| **Activity, Exploration** | **0.943** | **[0.872, 1.00]** |
| **Activity, Affiliation** | **-0.481** | **[-0.753, -0.219]** |
| **Activity, Aggression** | **0.623** | **[0.292, 0.956]** |
| Activity, Antennation | 0.00306 | [-0.292 0.301] |
| **Exploration, Affiliation** | **-0.352** | **[-0.599, -0.0413]** |
| **Exploration, Aggression** | **0.462** | **[0.108, 0.865]** |
| Exploration, Antennation | 0.00259 | [-0.295, 0.289] |
| Affiliation, Aggression | -0.231 | [-0.644, 0.242] |
| Affiliation, Antennation | -0.176 | [-0.482 0.127] |
| Aggression, Antennation | 0.104 | [-0.358, 0.533] |

**Figure 3:**
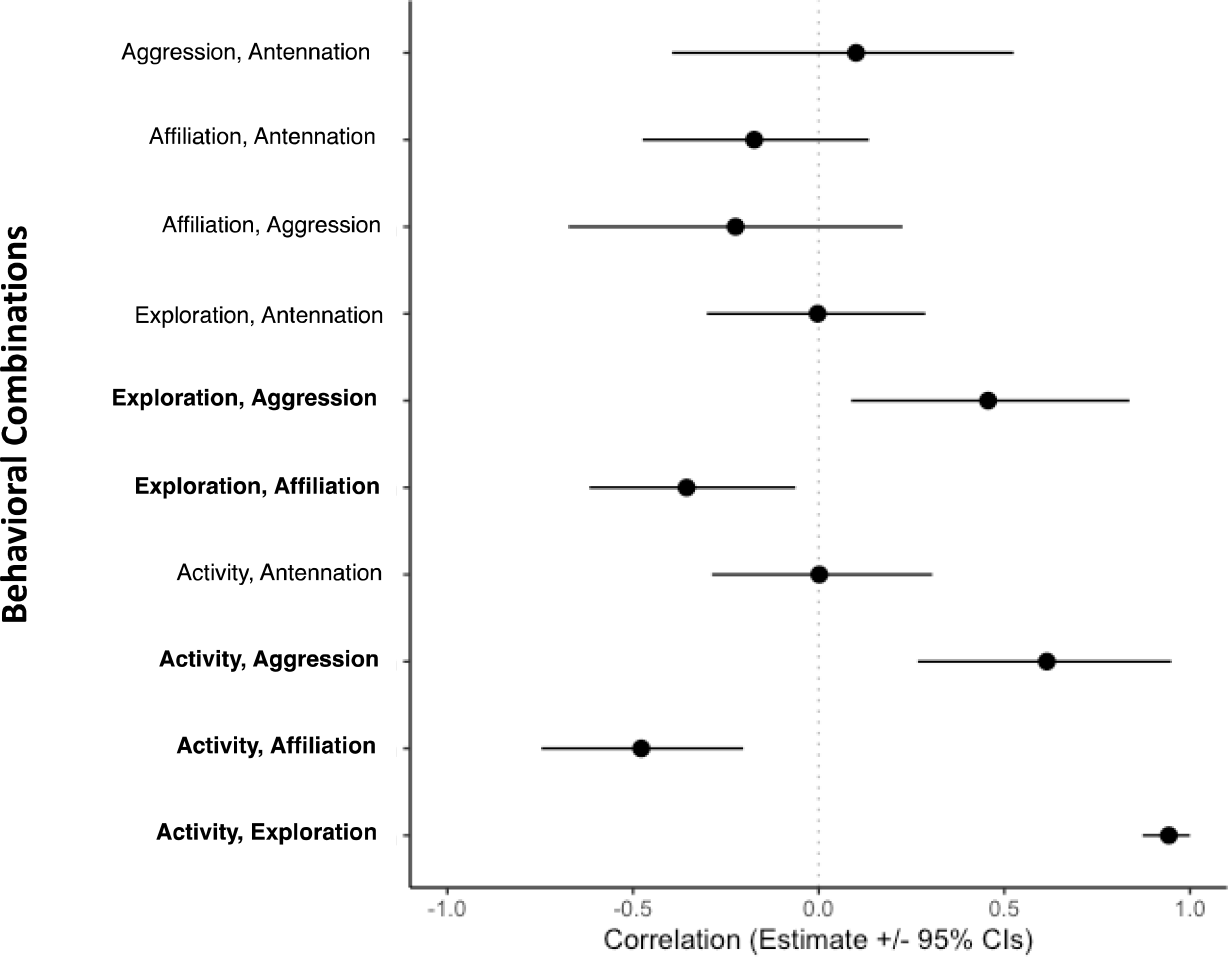
Correlation estimates from bivariate analysis. Five combinations of behaviors demonstrated significant correlations (bolded): Activity and Exploration (mean correlation: 0.943, 95% CI: [0.872, 1.00]), Activity and Aggression (mean correlation: mean correlation: 0.623, 95% CI: [0.292, 0.956]), Activity and Affiliation (mean correlation: -0.481, 95% CI: [-0.753, -0.219]), Exploration and Affiliation (mean correlation: -0.352, 95% CI: [-0.599, -0.0413]), and Exploration and Aggression (mean correlation: 0.462, 95% CI: [0.108, 0.865]).

## Discussion

Our study finds evidence of five different personality traits in *P. fuscatus*, and evidence of behavioral syndromes that encompass both social and non-social personality traits. Most excitingly, with two different methods of analysis we find significant correlations between both activity and aggression and activity and affiliation, but no correlations between antennation and any other personality trait. Correlations between non-social and social personality traits are particularly interesting because such links may play a role in the evolution and maintenance of variation in social phenotypes (Gartland et al. 2022; Laskowski et al. 2022).

Previous work has found a range of different types of relationships between social and non-social personality traits. We found a negative relationship between affiliative behavior and activity and a positive relationship between activity and aggression, suggesting that more active individuals engage in fewer affiliative and more antagonistic behavior with conspecifics. The positive relationship between aggression and activity found in this study is consistent with other work in sticklebacks (*Gasterosteus aculeatus)* (Dingemanse et al. 2007), chimpanzees (*Pan troglodytes*) (Koski 2011), red squirrels (*Tamiasciurus hudsonicus*) (Taylor et al. 2012), and crickets (*Gryllus integer*) (Kortet and Hedrick 2007). However, our findings contrast with previous studies that found positive correlations between activity and sociability in mammals (Petelle, Martin & Blumstein, 2015), reptiles (Michelangeli, Chapple & Wong, 2016), and fish (Cote et al. 2010). Activity is sometimes thought to be positively associated with sociability as active individuals may encounter more conspecifics than less active individuals (Petelle, Martin, and Blumstein 2015). However, social encounters may produce affiliative, aggressive, or neutral interactions. As a result, increased encounter rates may lead to more interactions, but not necessarily increased sociability. Additional work in other taxa will be important to assess links between activity and different types of social interactions. Although we find significant relationships between Activity and Affiliation and Activity and Aggression with both statistical analyses, we also find significant relationships between Exploration and Aggression and Exploration and Affiliation only when using a bivariate analysis. This finding is interesting as it suggests that behavioral syndromes may only be detected when also accounting for intra-individual covariance in behavior (Houslay and Wilson 2017; Houslay et al. 2018).

The evolutionary factors that maintain consistent individual variation in sociability remain controversial, with studies find conflicting fitness consequences for sociability (Silk et al. 2010; Yang, Maldonado-Chaparro, and Blumstein 2017). Recent work has theorized that individual differences in social personality can evolve and be maintained through the pace-of-life syndrome (POLS), where individuals face life-history trade-offs between maximizing immediate reproductive opportunities and survivorship, with bolder/more aggressive individuals engaging in more risky conflict to maximize short term reproductive gains (Gartland et al. 2022; Hall et al. 2015; Réale et al. 2010; Royauté et al. 2018; Wolf et al. 2007; Wolf, Sander Van Doorn, and Weissing 2008). Although little work has empirically investigated life-history trade-offs and social personality, our work suggests that social personalities may be subjected to trade-offs through linkage with other behavioral traits (Kim and Velando 2016). As we find a positive relationship between activity and aggression, but a negative relationship between activity and affiliation, our work suggests that there may be a “slow and social” phenotype in contrast to a “fast and aggressive” phenotype. Future work is needed to understand how these behavioral syndromes may or may not translate to life-history trade-offs in species with one reproductive season.

Our findings continue to support the theoretical and experimental linkage between activity and exploratory behavior. The strong positive correlation between activity and exploration we identified in wasps is consistent with that found in many vertebrate species such as fish (Budaev 1997; Cote et al. 2010), mammals, and birds (Hall et al. 2015). Previous invertebrate work has also found correlations between activity and exploration (Monceau et al. 2015). Although recent work finds that the strength of the links between activity and exploration may be influenced by the methods used to assess these traits, shared mechanisms are likely to explain some of the linkage of these two traits (Garamszegi, Markó, and Herczeg 2013). As is the case with many other studies (Koenig and Ousterhout 2018; Wilson and Godin 2009; Yuen et al. 2016), we use the same assay to evaluate both activity and exploration, which may contribute to the strong link between the two behaviors. However, the arena is large enough that wasps could move without entering new chambers, so wasps could be highly active without being exploratory. Similarly, wasps could take either more or less direct routes between chambers such that highly exploratory wasps may not be highly active. There is current interest in evaluating how genetics (Oers and Mueller 2010), physiological condition (Wu and Seebacher 2022), and hormonal pleiotropy (Stöwe et al. 2010) maintain linkage between activity and exploration. Further work is needed to understand if the same mechanisms that support linkage of activity and exploration drive relationships between other personality traits and when behavioral assays constrain this relationship (Oers and Mueller 2010).

Interestingly, we find that there is very little difference between repeatability calculated with additional individual and trial factors included in models, with fixed effects accounting for relatively small proportion of variation in behavior, with the exception of Aggression. Trial effects (trial order and dummy ID) and body mass account for more variation in Aggression than individual ID, indicating that individuals are more likely to alter aggressive behavior than affiliative behavior or antennation in response to dummy wasps. While some work predicts that behaviors that depend on social context should be most variable, our results contrast with one of the largest meta-analyses to date which finds aggressive behavior to be one of the most repeatable across taxa (Bell, Hankison, and Laskowski 2009). The variation in Aggression due to trial conditions we observe is consistent with relative resource holding potential altering contest dynamics in other *polistine* wasp species (Cini et al. 2011; Tibbetts and Shorter 2009). Future work could consider comparing repeatability of aggressive behavior to size-matched and non-sized matched dummies.

We find very different repeatability of personality traits compared to the one other study evaluating personality traits in *Polistes* wasps (Wright et al. 2018). Wright et al. (2018) assessed aggression, exploration, and activity in *Polistes metricus* wasps, finding much higher repeatability than this study (aggression Wright et al.: R = 0.88, aggression Jomaa et al. “simple” R = 0.141, aggression Jomaa et al. “adjusted” R: 0.159; exploration Wright et al.: R = 0.88, exploration Jomaa et al. “simple”: R = 0.497, exploration Jomaa et al. “adjusted” R: 0.513; activity Wright et al.: R = 0.92, activity Jomaa et al. “simple”: R = 0.378, activity Jomaa et al. “adjusted”: R = 0.398). Our aggression assay differed from that used by Wright, as we assessed social aggression and Wright focused on defensive aggression. However, the assay used for exploration and aggression was similar in the two studies. We originally thought Wright et al. (2018) may have high repeatability because they included multiple additional factors in the models: head size, starting nest size, egg count, and trial number as fixed effects, and wasp ID nested within site ID and dummy wasp ID as random effects. However, including additional fixed effects in our model (wasp body weight, trial number, and dummy ID) did not substantially increase repeatability. Repeatability values reported in Wright et al. (2018) are significantly higher than typical values reported in other personality studies for Hymenoptera (Gomes, Desouhant, and Amat 2019; Monceau et al. 2015) and other species such as mammals (Wat, Banks, and McArthur 2020), fish (Jones and Godin 2010), and birds (Dingemanse et al. 2002). Inflated repeatability measures may lead to misleading conclusions about variation in populations and upper limits of repeatability of traits (Wilson 2018). Further work that explores how and why personality repeatability varies may be useful for understanding the differences across studies.

Eusocial insects are exciting models for examining the ontogeny, mechanisms, and fitness consequences of behavior at multiple levels of social organization (Jandt et al. 2014). Work in other social insects has revealed personality traits both within and between colonies for many behaviors including aggression (Suarez et al. 2002), exploration (Modlmeier and Foitzik 2011), and cooperation (Robinson, Page, and Fondrk 1990). Perhaps most interestingly, work has begun to examine how colony behavioral phenotypes provide fitness benefits in different environmental conditions (Bengston and Dornhaus 2015; Blight et al. 2016; 2017; Segev et al. 2017). Previous work in *Polistes* has examined the relationship between queen personality and colony behavior, and found that queen personality negatively predicts colony response to attack (Wright et al. 2017). In contrast to most eusocial insect societies, *P. fuscatus* colonies are singly or multiply founded with no morphological caste differences between queens and workers, providing an opportunity to study how behavioral syndromes influence colony foundation, dominance hierarchies, and cooperation. Furthermore, as an emerging model system for neurobiology, *P. fuscatus* present an excellent system to examine neural and genomic linkages in behavioral syndromes (Berens et al. 2015).

Our work demonstrates that *P. fuscatus* wasps have repeatable personality traits and behavioral syndromes, including consistent individual variation in sociability. A growing body of evidence illustrates that consistent individual variation in social and non-social personality can have broadly important ecological, evolutionary, and behavioral effects (Sih et al. 2012; Laskowski et al. 2022). Moreover, understanding how social behavior traits are correlated with non-social traits provides insight into how variation in social phenotypes is maintained within populations. *P. fuscatus* have potential to add to this work as a facultatively eusocial model taxa for exploring the causes and consequences of individual behavioral variation.

## Acknowledgements

Wasps were collected with assistance from Zachary Leytus. Data collection was performed by Katie Mclean and Fatima Jomaa. Wasp personality assays were conducted by Fatima Jomaa, Anna Vi, Micah Golan, and Fiona Corcoran. This work was supported by the University of Michigan, National Geographic Society, and the National Science Foundation Grants IOS 1557564 and 2134910.

## Author Contributions

Fatima W. Jomaa collected and analyzed data and wrote initial drafts of the manuscript. Emily C. Laub designed assays, analyzed data, and wrote the manuscript. Elizabeth A. Tibbetts designed the experiment and provided crucial insights and edits to the manuscript.

